# Regulation of oncogene-induced senescence by the MRE11 and TREX1 nucleases

**DOI:** 10.1101/2023.03.30.534897

**Authors:** Hervé Técher, Diyavarshini Gopaul, Jonathan Heuzé, Baptiste Leray, Audrey Vernet, Clément Mettling, Jérôme Moreaux, Yea-Lih Lin, Philippe Pasero

## Abstract

Oncogene-induced senescence (OIS) is a tumor-suppressive mechanism that arrests cell proliferation in response to oncogene-induced replication stress (RS). OIS also depends on the cGAS-STING pathway, which detects cytosolic DNA and promotes the expression of type I interferons (IFN) and pro-inflammatory cytokines. Whether and how the RS and IFN responses cooperate to promote OIS is currently unknown. Here, we show that the MRE11 nuclease promotes OIS in immortalized human fibroblasts overexpressing the H-RAS^V12^ oncogene both by slowing replication forks and by activating the cGAS-STING pathway in response to RS. Interestingly, overexpression of TREX1, the major nuclease degrading cytosolic DNA, prevented RAS-induced senescence. In contrast, overexpression of a dominant negative mutant of TREX1 was sufficient to induce senescence in human fibroblasts, even in the absence of H-RAS^V12^ induction. Collectively, these data suggest that the RS and IFN responses in OIS are functionally linked through a process involving the nucleases MRE11 and TREX1.

## Introduction

Oncogene-induced senescence (OIS) is a process in which cells stop proliferating due to genomic instability caused by activated oncogenes (Gorgoulis *et al*, 2019; Kotsantis *et al*, 2018). As such, OIS represents a major barrier to cell transformation, but how this process is initiated at the molecular level is currently unclear. Two non-exclusive mechanisms have been proposed. Alteration of the DNA replication program by deregulated oncogenes has been shown to generate DNA damage and induce senescence through activation of the DNA damage response (DDR) (Bartkova *et al*, 2006; Di Micco *et al*, 2006). In addition, recent evidence indicates that OIS also depends on the cyclic GMP-AMP Synthase (cGAS) and the stimulator of interferon genes (STING) pathway (Yang *et al*, 2017; Glück *et al*, 2017; Dou *et al*, 2017). It is currently unknown whether the DDR and cGAS-STING pathways cooperate to induce OIS or act independently.

Oncogene-induced replication stress is characterized by alterations in fork progression and accumulation of DNA damage and chromosomal aberrations (Bartkova *et al*, 2006, 2005; Di Micco *et al*, 2006). One of the best-characterized examples of OIS is caused by the overexpression of H-RAS^V12^, which leads to the constitutive activation of the MAPK/ERK pathway and drives cell transformation. In immortalized human fibroblasts, H-RAS^v12^ deregulates dNTP pools and increases transcription-replication conflicts, leading to increased genomic instability (Kotsantis *et al*, 2016; Maya-Mendoza *et al*, 2015; Di Micco *et al*, 2006; Bianco *et al*, 2019). We have recently shown that immortalized human BJ fibroblasts overexpressing H-RAS^V12^ (BJ-RAS^V12^) escape senescence by overexpressing Claspin and Timeless, two components of the fork protection complex (Bianco *et al*, 2019). This overexpression, which was also observed in primary breast, lung and colon cancers, allowed BJ-RAS^V12^ cells to escape RS and restore a normal fork progression, supporting the view that RS plays a central role in OIS (Bianco *et al*, 2019; Kotsantis *et al*, 2016).

The cytosolic DNA sensor cGAS recognizes double-stranded DNA (dsDNA) species longer than 40 bp and produces cyclic guanosine monophosphate–adenosine monophosphate (cGAMP) as a second messenger to activate STING (Ablasser & Chen, 2019). Then, the STING-TBK1-IRF3 axis activates the transcription of type-I interferon genes (IFN-α and IFN-β genes) that subsequently promotes the expression of interferon-stimulated genes (ISGs).

Altogether, these pro-inflammatory factors, once produced and secreted, act as immuno-modulators. Senescence is associated with the secretion of cytokines and chemokines through a process known as the senescence-associated secretory phenotype (SASP) (Coppé *et al*, 2008). Overexpression of the major cytosolic 3’-exonuclease TREX1 decreases the level of SASP during OIS (Takahashi *et al*, 2018), suggesting that the level of cytosolic DNA determines the SASP response. Although SASP is mediated by the canonical cGAS-STING pathway (Dou *et al*, 2017; Glück *et al*, 2017), cell proliferation arrest could also depend on cGAS in a STING-independent manner (Yang *et al*, 2017; Dou *et al*, 2017). This apparent discrepancy raises the question of the origin of cytosolic DNA and the mechanism by which it contributes to senescence. It has been observed that cells overexpressing oncogenes suffer from mitotic failures (Dikovskaya *et al*, 2015) and accumulate micronuclei (Dou *et al*, 2017; Kotsantis *et al*, 2016). Since cGAS binds DNA inside damaged micronuclei, cytosolic DNA could provide the signal for cGAS-mediated senescence and SASP (Dou *et al*, 2017; Glück *et al*, 2017).

Here, we used immortalized BJ-hTERT fibroblasts overexpressing the H-RAS^V12^ oncogene under a tetracycline-inducible promoter to investigate the molecular links between oncogene-induced RS and cGAS-mediated DNA sensing in the establishment of OIS. We show that the inhibition of MRE11 by Mirin abrogated RAS^V12^-induced senescence, fork progression defect and micronuclei formation in BJ fibroblasts. Furthermore, overexpression of TREX1 or inhibition of cGAS in BJ-RAS^V12^ cells decreased the percentage of senescent cells. In contrast, the expression of a dominant-negative form of TREX1 (D18N) or supplementation with interferon-β (IFN-β) induced senescence even in the absence of H-RAS^V12^ induction. Collectively, our results show that the nucleases MRE11 and TREX1 control OIS by modulating RS and cytosolic DNA sensing. Our work reveals a crosstalk between the canonical RS response and the cGAS-STING pathway at the onset of OIS, which has important implications for cancer biology.

## Results

### RAS-induced replication stress correlates with the induction of IFN, ISG and SASP genes

Oncogenes such as H-RAS^V12^ induce senescence by causing replication stress and activating DDR signaling (Gorgoulis *et al*, 2019). Recent studies indicate that OIS depends also on the cGAS-STING pathway (Glück *et al*, 2017; Dou *et al*, 2017; Yang *et al*, 2017), but whether and how the DDR and cGAS-STING pathways cooperate to induce OIS is currently unknown. To address these questions, we first monitored the expression of IFN, ISG and SASP genes before and after OIS. To this end, oncogenic stress was induced in telomerase-immortalized human BJ fibroblasts expressing the H-RAS^V12^ gene under the control of a doxycycline-inducible promoter and differential gene expression was analyzed by RNA-seq as described previously (Bianco *et al*, 2019). Volcano plots revealed an increased expression of IFN, ISG and SASP genes in BJ-RAS^V12^ cells compared to control BJ cells (**Fig. 1A**), which is consistent with earlier studies (Glück *et al*, 2017; Dou *et al*, 2017; Yang *et al*, 2017).

**Figure 1.**
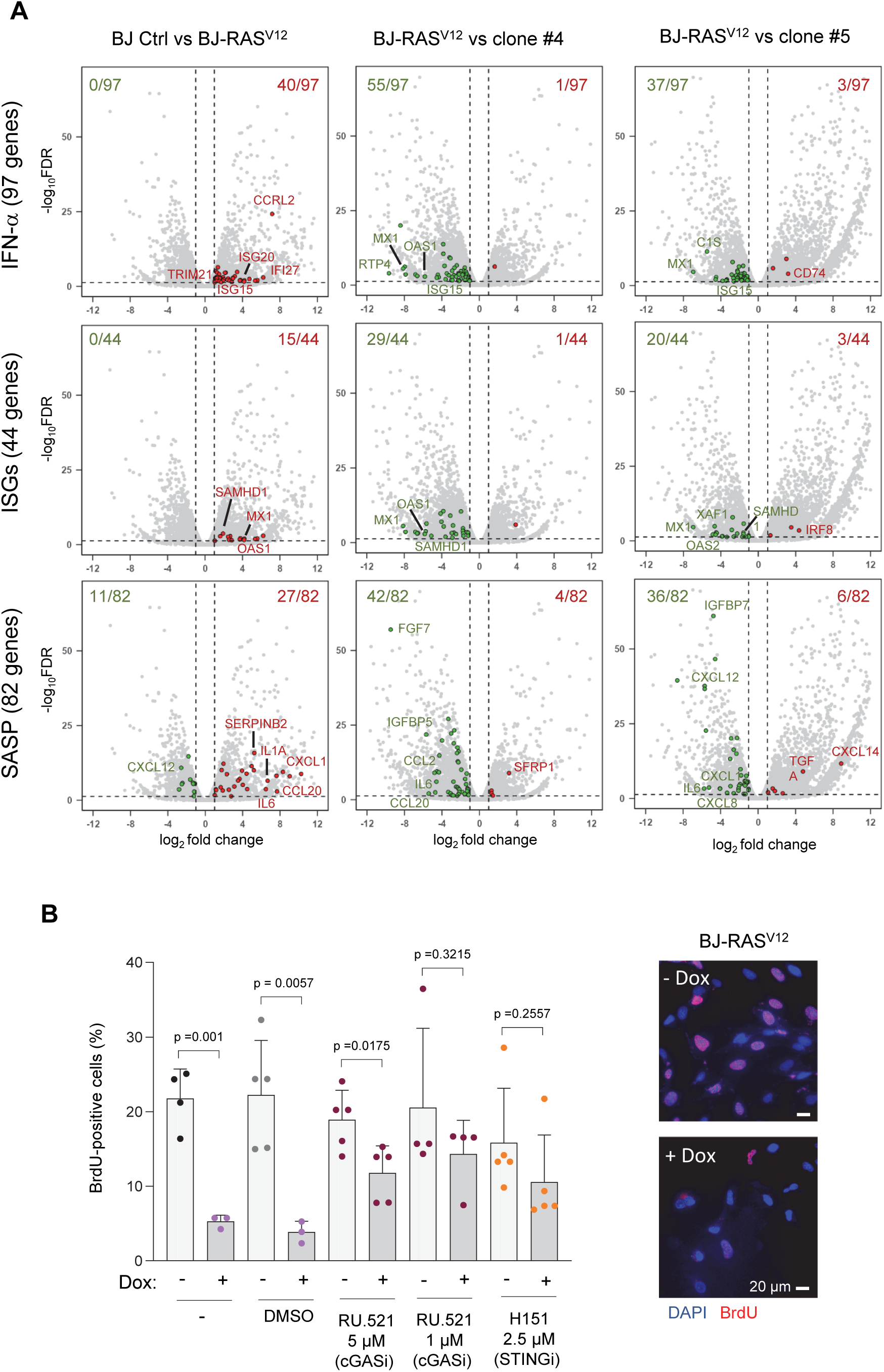
Induction of IFN, ISG and SASP genes in response to RAS^V12^-induced RS. (**A**) Volcano plots of differentially expressed genes in BJ-RAS^V12^ cells expressing or not RASV^12^ and in senescent BJ-RAS^V12^ cells escaping senescence by overexpressing Claspin and Timeless (clones #4 and #5; Bianco et al, 2019). Gene sets correspond to interferon-α response genes (IFN-α, GSEA M5911), interferon stimulated genes (ISGs, Reactome R-HSA-9034125) and SASP genes (from (Ruiz-Vega *et al*, 2020)). Genes that are significantly up-regulated are shown in red (log_2_ fold change > 1 and FDR < 0.05). Genes significantly down-regulated are shown in green (log_2_ fold change < -1 and FDR < 0.05). Data are from biological duplicates. (**B**) The cGAS-STING pathway contributes to the cell proliferation arrest induced by RAS^V12^ in BJ fibroblasts. The overexpression of RAS^V12^ was induced (+10 μg/ml Dox) or not (-Dox) for 8 days and cells were pulse-labeled with 30 μM BrdU for 20 minutes. BrdU incorporation was monitored by immunofluorescence (see methods section). To determine the role of cGAS and STING in cell proliferation arrest, cells were treated for the same period of time with the cGAS inhibitor RU.521 (1 or 5 μM) and the STING inhibitor H151 (2.5 μM). DMSO was used as a control. The frequency of BrdU-positive cells is shown. Mean, standard deviation (SD) and p-values (unpaired t-test) are shown for at least three independent experiments. Representative images of induced (+Dox) and non-induced (-Dox) BJ-RAS^V12^ fibroblasts are shown. BrdU: red, DAPI: blue. Scale bars are 20 μm.

We have recently shown that BJ-RAS^V12^ cells adapt to oncogene-induced RS and escape OIS by overexpressing downstream components of the ATR-CHK1 pathway such as Claspin and Timeless (Bianco *et al*, 2019). To determine whether this adaptation affects the expression of IFN, ISG and SASP genes, we analyzed gene expression by RNA-seq in the clones that escaped senescence. Remarkably, two clones that escaped OIS by overexpressing Claspin and Timeless (clones #4 and #5) showed a marked reduction in IFN, ISG and SASP gene expression compared to senescent BJ-RAS^V12^ fibroblasts (**Fig. 1A**). In contrast, the clone #8 that escaped OIS independently of Claspin and Timeless overexpression (Bianco *et al*, 2019) did not display changes in the expression of IFN and ISG genes, and a reduction of SASP genes relative to BJ-RAS^V12^ cells (**EV Fig. 1**).

To confirm that activation of the cGAS-STING pathway contributes to senescence in BJ-RAS^V12^ fibroblasts, we next evaluated the effect of cGAS-STING pathway inhibitors on cell proliferation arrest, an early step in OIS. To this end, we measured BrdU incorporation in BJ-RAS^V12^ cells treated or not for 8 days with small molecule inhibitors targeting either cGAS (RU.521, 1 or 5 μM) or STING (H151, 2.5 μM) (Haag *et al*, 2018; Wiser *et al*, 2020; Vincent *et al*, 2017). RAS^V12^ induction reduced the frequency of BrdU-positive cells by 76 to 83% relative to untreated cells. However, this reduction was only of 31 to 38% when BJ-RAS^V12^ cells were treated with the cGAS and STING inhibitors (**Fig. 1B**), which is consistent with previous reports (Yang *et al*, 2017; Glück *et al*, 2017). Collectively, these results indicate that oncogene-induced replication stress and activation of the cGAS-STING pathway are functionally linked during RAS^V12^-induced senescence in BJ fibroblasts.

### MRE11 contributes to the IFN response and SASP in BJ-RAS^V12^ cells

The recovery of stalled replication forks depends on the processing of nascent DNA by a variety of DNA helicases and nucleases (Pasero & Vindigni, 2017). Central to this process is MRE11, an exo/endonuclease that initiates the resection of nascent DNA to promote HR-mediated fork restart (Petermann *et al*, 2010; Hashimoto *et al*, 2012; Haince *et al*, 2008; Bryant *et al*, 2009). Since MRE11 is required for the accumulation of cytosolic DNA in response to RS (Coquel *et al*, 2018), we next asked whether MRE11 could also mediate the oncogenic activation of the cGAS pathway. To address this possibility, we analyzed the induction of the IFN response by RNA-seq in BJ-RAS^V12^ fibroblasts treated or not with the MRE11 inhibitor Mirin (Dupré *et al*, 2008). This analysis revealed a reduction of IFN, ISG and SASP gene expression after treatment with 10 μM Mirin for 8 days (**Fig. 2A**), which was confirmed by RT-qPCR for the IL6 and CXCL1 genes (**Fig. 2B**) and by immunoblotting for IL-1α (**EV Fig. 2**). We also found that the reduction in gene expression induced by Mirin treatment was not due to the alteration in the subcellular localization of MRE11 and cGAS upon H-RAS^V12^ induction (**EV Fig. 2**). These data show that MRE11 contributes to the activation of the IFN response in BJ-RAS^V12^ fibroblasts.

**Figure 2.**
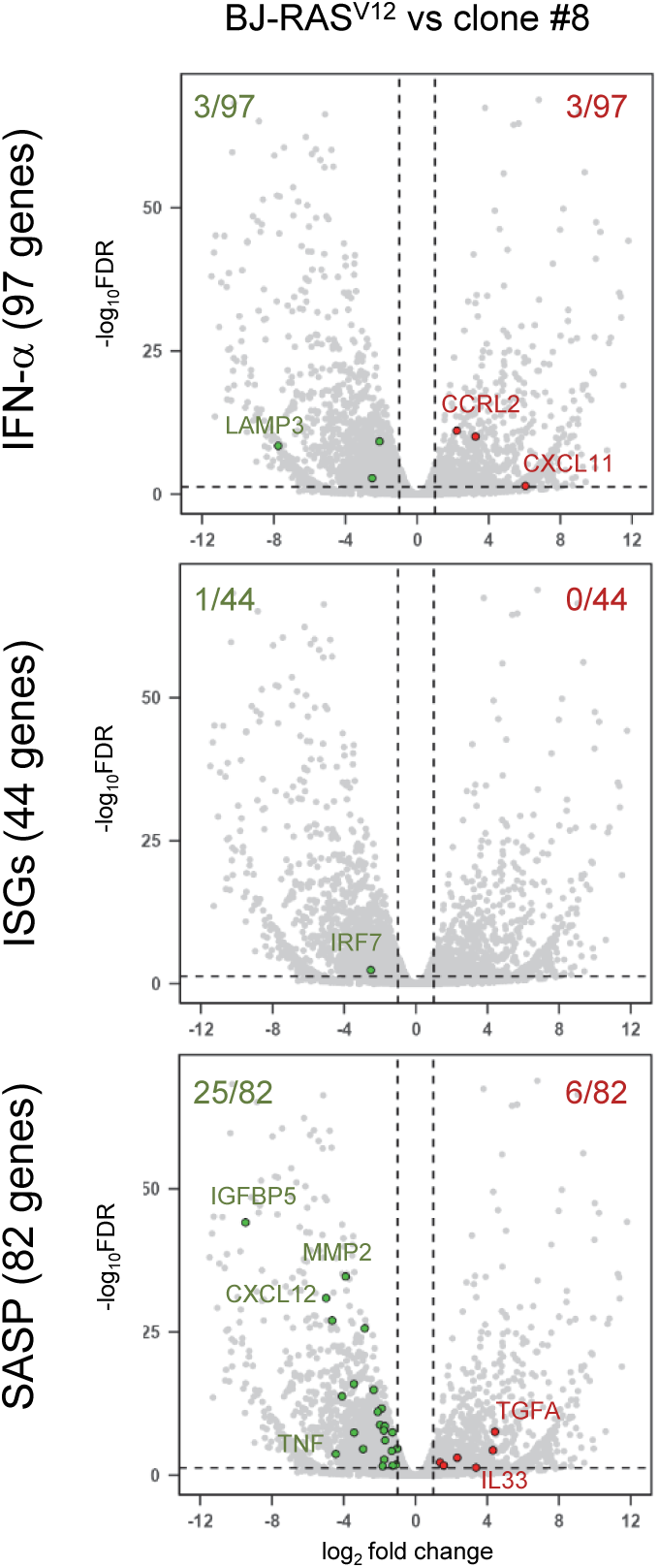
The induction of IFN, ISG and SASP genes in BJ-RAS^V12^ cells depends on MRE11. (**A**) BJ-RAS^v12^ fibroblasts were induced for 8 days with 10 μg/ml Dox in the presence or the absence of 10 μM Mirin. Volcano plots of differentially expressed IFN, ISG and SASP genes between control and RAS-induced BJ cells treated or not with Mirin are shown. RNA-seq data were processed as described in Fig. 1A. Results correspond to biological duplicates. (**B**) Levels of IL6 and CXCL1 RNA were quantified by RT-qPCR to confirm RNA-seq data.

### MRE11 is required for OIS in BJ-RAS^V12^ cells

To further investigate the role of MRE11 in OIS, we monitored the effect of Mirin on cell proliferation after H-RAS^V12^ induction. As shown in **Figure 3A**, H-RAS^V12^ strongly reduced the proliferation of BJ cells after two passages (∼8 days post induction). However, this growth inhibition was completely suppressed when cells were treated with 10 µM Mirin (**Fig. 3A**), indicating that MRE11 plays a major role in the growth arrest induced by RAS^V12^.

**Figure 3.**
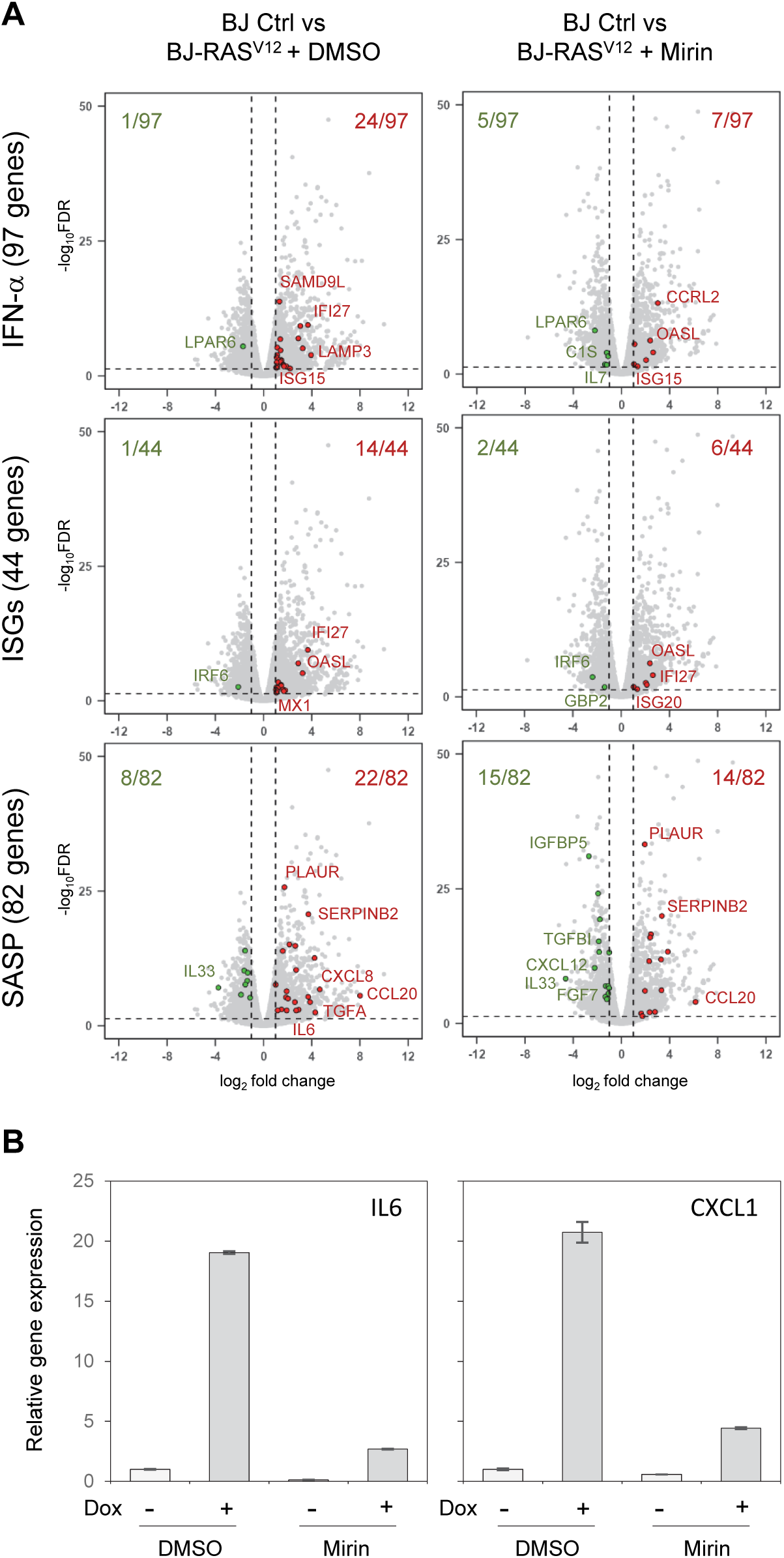
MRE11 is essential for the induction of senescence in BJ-RAS^V12^ fibroblasts. (**A**) Effect of MRE11 inhibition on the proliferation of BJ-RAS^V12^ cells. Untreated (Ctrl) or RAS-induced (Dox,10 μg/ml) BJ-RAS^V12^ cells were grown in the presence (Mirin, 10 µM) or the absence (DMSO) of the MRE11 inhibitor Mirin. Cumulative cell number was calculated by counting cells during five consecutive passages, as indicated Materials and methods. One representative experiment is shown (n=2). (**B**) MRE11 inhibition prevents the induction of senescence-associated β-galactosidase (SA-β-gal). BJ-RAS^v12^ fibroblasts induced with Dox (10 μg/ml) for 8 and 14 days in the presence or the absence of 10 μM Mirin were stained for SA-β-gal activity. The frequency of SA-β-gal positive cells was scored by microscopy. Representative images are shown in EV Fig. 3A. Mean +/-SD (n=3) and p-values (unpaired t-test) are shown. (**C**) Differential effect of MRE11 inhibitors on cell proliferation. The frequency of BrdU-positive cells before and after RAS^V12^ induction was monitored as described in Fig. 1B. Cells were either treated with Mirin or with two Mirin derivatives, PFM01 and PFM39, which target the endonuclease or the exonuclease activities of MRE11, respectively. All compounds were all used at a final concentration of 10 μM. Non-treated (-) and DMSO-treated cells were used as controls. Mean, SD and p-values (unpaired t-test) are shown for at least two independent experiments.

Next, we monitored the effect of MRE11 inhibition on the induction of senescence-associated β-galactosidase (SA-β-gal), a hallmark of senescent cells. The percentage of BJ-RAS^V12^ fibroblasts exhibiting β-galactosidase activity increased significantly on day 8 and day 14 after H-RAS^V12^ induction. However, this SA-β-gal activity was abolished after Mirin treatment (**Fig. 3B**), consistent with the high proliferative activity of these cells (**Fig. 3A**). Mirin also restored BrdU incorporation in BJ-RAS^V12^ cells (**Fig. 3C** and **EV Fig. 3**) to the same extent as cGAS and STING inhibitors (**Fig. 1B**), supporting the view that MRE11 plays an important role in OIS.

Mirin is a potent inhibitor of the exonuclease activity of MRE11 (Dupré *et al*, 2008), but this enzyme also has an endonuclease activity. To determine whether this activity is also involved in OIS, BJ-RAS^V12^ cells were treated with inhibitors differentially targeting the endonuclease (PFM01) or the exonuclease (PFM39) activities of MRE11 (Shibata *et al*, 2014), all used at the same concentration here (10 µM). The fraction of cells incorporating BrdU after 8 days of H-RAS^V12^ induction was restored by PFM01 to the same extent as Mirin, but was not restored as efficiently by PFM39 (**Fig. 3C**). Together, these data suggest that both the endonuclease and exonuclease activities of MRE11 contribute to OIS.

### MRE11 promotes oncogene-induced replication stress in BJ-RAS^V12^ cells

Our results indicate that MRE11 plays an active role in the induction of the IFN pathway and in OIS. To determine whether MRE11 acts upstream of these pathways by contributing to oncogene-induced RS, we measured fork progression by DNA fiber spreading in BJ-RAS^V12^ fibroblasts treated or not with 10 μM Mirin (**Fig. 4A** and **EV Fig. 4A**). As previously reported (Kotsantis *et al*, 2016; Bianco *et al*, 2019), we observed a significant reduction in fork speed upon H-RAS^V12^ induction for 8 days (**Fig. 4A** and **EV Fig. 4B**), indicative of oncogene-induced RS. Remarkably, normal fork velocity was restored when cells were treated with Mirin either immediately upon H-RAS^V12^ induction (**Fig. 4A**) or five days post induction (**EV Fig. 4C**). These results are consistent with the fact that Mirin restored BrdU incorporation under the same conditions (**Fig. 2C** **and EV** **Fig. 3B**). We also observed an increased frequency of asymmetric IdU and CldU track lengths in BJ-RAS^V12^ cells, indicative of RAS-induced fork arrest (Técher *et al*, 2013). This fork asymmetry was reduced in the presence of Mirin (**Fig. 4B**). To determine whether MRE11 also contributes to the oncogene-induced DNA damage response, we assessed the phosphorylation of DDR factors by western blotting 8 days after RAS^V12^ induction. As expected, RAS^V12^ increased the levels of phospho-ATM (S1981) and phospho-RPA32 (S4/S8 and S33) in BJ-RAS^V12^ fibroblasts. This DDR induction was also prevented by Mirin (**Fig. 4C**), which is consistent with earlier studies (Dupré *et al*, 2008). Of note, p53 protein level was not affected by H-RAS^V12^ expression or Mirin treatment (**Fig. 4C**). Taken together, these results indicate that the MRE11 nuclease enhances oncogene-induced replication stress in BJ-RAS^V12^ cells and contributes to the activation of the DDR pathway.

**Figure 4.**
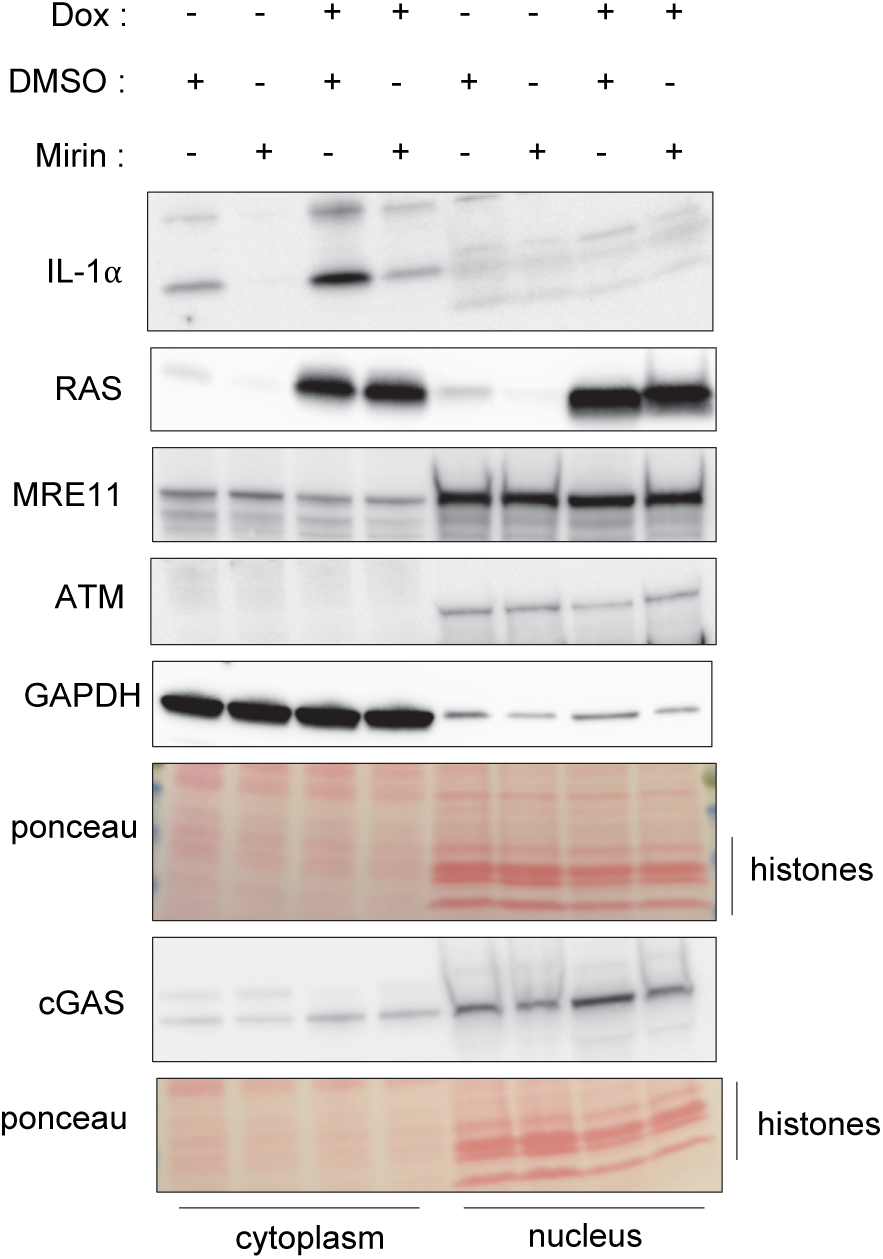
Mirin prevents RAS-induced replication stress. (**A**) BJ-RAS^V12^ cells were grown for 8 days with or without Dox, DMSO and Mirin as indicated. Cells were labeled with two consecutive pulses of IdU and CldU for 20 minutes each and DNA fibers were stretched on glass slides as indicated in Materials and Methods. The length of IdU and CldU tracks, indicative of fork speed, was measured for at least 200 individual ongoing forks (adjacent red and green tracks; see EV Fig. 4A). Results are displayed as super-plots of the distribution of IdU and CldU track length (μm) for 2-3 biological replicates. Median and P-values (two-sided paired-t test) are indicated. (**B**) Mirin suppresses the increased fork asymmetry induced by RAS^V12^. Fork asymmetry, which is indicative of increased fork pausing or stalling, was measured as the ratio of the longest to the shortest track for each individual fork in BJ-RAS^V12^ cells treated with different combinations of Dox, DMSO and Mirin. Increased fork asymmetry in RAS-induced cells, indicative of replication stress, is suppressed by MRE11 inhibition. Median and p-values (non-parametric Mann-Whitney rank sum test) are indicated. A representative experiment is shown (n=2). (**C**) Western blot analysis of DDR factors in BJ cells overexpressing (+Dox) or not (-Dox) RAS^v12^ and treated or not with 10 µM Mirin. Cytosolic and nuclear fractions were obtained by lysing cells for 5 minutes on ice with a lysis buffer (10 mM Tris pH7.4, 10 mM NaCl, 10 mM MgCl_2_) containing 0.25% NP-40. Nuclear pellets were lysed in 2x Laemmli buffer and digested with benzonase. (**D**) Mirin prevents the accumulation of micronuclei in BJ-RAS^V12^ fibroblasts. The frequency of micronuclei was determined in cells treated with various combinations of Dox and Mirin, as indicated in panel A. Mean, SD and p-values (unpaired t-test) are shown for 3 independent experiments.

### Mirin prevents the formation of micronuclei in BJ-RAS^V12^ cells

Micronuclei accumulate frequently under mild replication stress conditions as a consequence of unrepaired DSBs, incomplete DNA replication, and chromosome non-disjunction (Wilhelm *et al*, 2019; Chan *et al*, 2009). Micronuclei have been implicated in the activation of the cGAS-STING pathway through rupture of their nuclear envelope (Harding *et al*, 2017; Mohr *et al*, 2021; Mackenzie *et al*, 2017). Since MRE11 plays an active role in RAS-induced RS (**Fig. 4A** and **4B**), we then asked whether it could activate the cGAS-STING pathway through the formation of micronuclei. In BJ-RAS^V12^ fibroblasts, the frequency of micronuclei increased by ≍10-fold after 5 days of RAS^V12^ induction and further increased after 8 days (**Fig. 4D** and **EV Fig. 4D**). To determine whether this increased frequency of micronuclei was dependent on MRE11, BJ-RAS^V12^ fibroblasts were treated with or without 10 μM Mirin in the presence or absence of RAS^V12^ induction. Mirin treatment did not affect the level of micronuclei in uninduced cells, while it dramatically attenuated micronuclei formation after 5 and 8 days of H-RAS^V12^ induction (**Fig. 4D**), indicating that MRE11 plays an active role in the formation of RAS-induced micronuclei.

### TREX1 activity fine tunes OIS

To assess the impact of micronuclei and cytosolic DNA on senescence, we next investigated the effect of overexpressing the major cytosolic nuclease TREX1 on OIS. To this end, we constructed BJ-RAS^V12^ cells constitutively expressing GFP-tagged versions of either the wild-type (TREX1) or a dominant-negative nuclease-dead (TREX1-D18N) protein (Lee-Kirsch *et al*, 2007; Lehtinen *et al*, 2008) (**Fig. 5A**). Both proteins were predominantly located in the cytosol (**EV Fig. 5A**) and accumulated within ∼35% of micronuclei (**EV Fig. 5B**). Overexpression of TREX1-GFP reduced the level of RAS-induced SA-β-gal activity to that of control cells (**Fig. 5B**), supporting the view that cytosolic DNA contributes to OIS. In contrast, overexpression of the nuclease-dead mutant TREX1-D18N did not attenuate OIS in cells overexpressing H-RAS^V12^ (**Fig. 5B**). Of note, overexpression of TREX1-D18N was sufficient to induce SA-β-gal in BJ fibroblasts, even in the absence of H-RAS^V12^ expression (**Fig. 5B**), which is consistent with its dominant negative effect on endogenous TREX1. Using RNA-seq, we found that the TREX1-D18N mutant increased the expression of type I-IFN signaling, ISG and SASP genes (**Fig. 5C** and **EV Fig. 5C**), which was confirmed by RT-qPCR for IL6 and CXCL1 (**Fig. 5D**). Together, these results suggest that TREX1 negatively regulates OIS by degrading cytosolic DNA. Conversely, nuclease-dead TREX1 is sufficient to trigger senescence independently of RAS^V12^ induction, presumably by allowing accumulation of cytosolic DNA.

**Figure 5.**
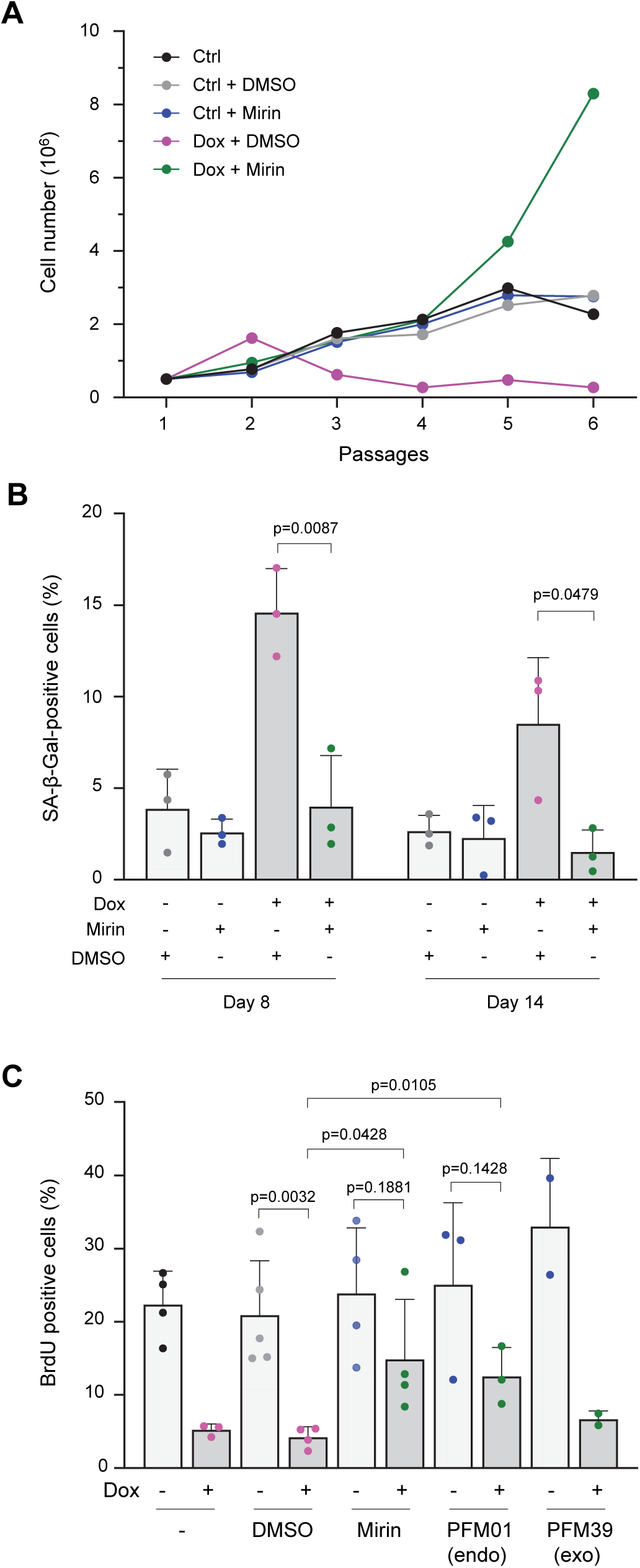
The TREX1 nuclease modulates senescence in BJ-RAS^V12^ fibroblasts. (**A**) Western blotting analysis of the levels of RAS^v12^ and TREX1 in BJ-RAS^V12^ cells that stably express the full-length (TREX1) or D18N mutant of TREX1 (TREX1-D18N) tagged with GFP. (**B**) Frequency of SA-β-gal positive cells in BJ-RAS^V12^ fibroblasts expressing TREX1 and TREX1-D18N. Mean, SD and p-values (unpaired t-test) are shown for three biological replicates. (**C**) RNA-seq analysis SASP and IFN-α response genes in TREX1 and TREX1-D18N overexpressing cells. RNA-seq samples were treated and analyzed as in Fig. 1A. Data are from two biological replicates and are compared to non-induced BJ-RAS cells (BJ-Ctrl). (**D**) IL6 and CXCL1 RNA levels in indicated cell lines, quantified by RT-qPCR.

### IFN-β induces replication stress and senescence independently of RAS^V12^

We next asked whether BJ-RAS^V12^ cells could be driven into senescence by the cGAS-STING pathway independently of RAS^V12^ induction. To address this possibility, BJ-RAS^V12^ cells were supplemented for 8 days with 300 U/ml of recombinant interferon-β (IFN-β) and the fraction of cells entering senescence was assessed using the SA-β-gal assay. Strikingly, IFN-β induced a higher percentage of SA-β-gal positive cells than RAS^V12^ (**Fig. 6A** and **EV Fig. 6A**). Consistently, IFN-β also reduced the frequency of BrdU-positive cells to the same level as that of RAS^V12^, even when used at a lower concentration (50 U/ml) (**Fig. 6B**). We also observed a mild increase in the expression of ISG15 when BJ-RAS^v12^ cells were exposed to 50 U/ml of IFN-β (**EV Fig. 6B**).

**Figure 6.**
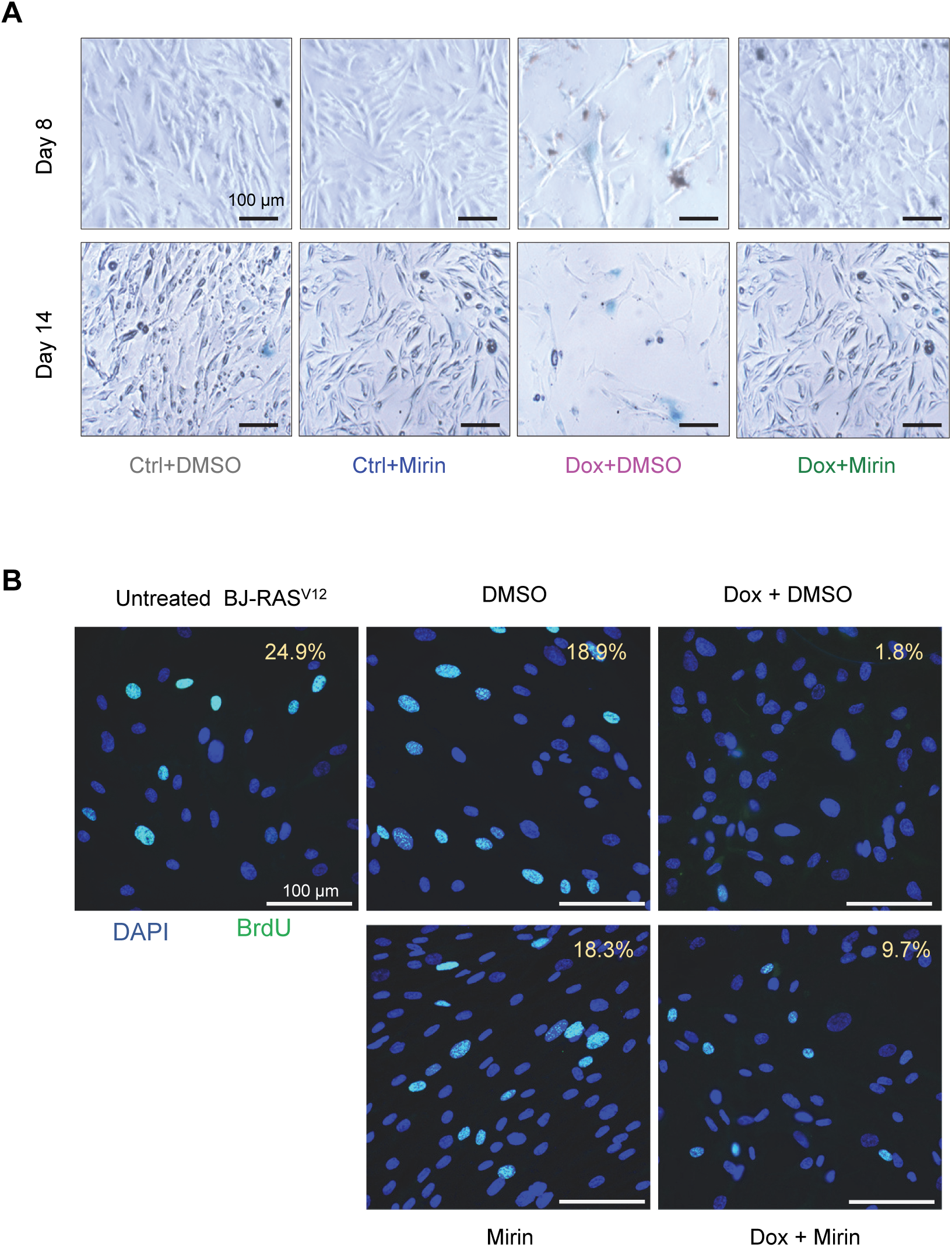
IFN-β induces replication stress and senescence in BJ fibroblasts. (**A**) BJ-RAS^V12^ fibroblasts (+/-Dox) were treated or not with 300 U/ml of recombinant interferon-β (IFN-β) for a period of 8 days and the frequency of SA-β-gal positive cells was scored in two independent experiments. (**B**) The proliferation of BJ-RAS^V12^ cells treated as in panel A was assessed by BrdU staining experiments. IFN-β was used at a final concentration of either 50 or 300 U/ml. Mean, SD and p-values (unpaired t-test) are shown fork at least two independent experiments. (**C**) DNA fiber analysis of fork progression in the indicated cells and conditions. Analysis was performed as described in Fig. 3A. (**D**) Micronuclei frequency in BJ-RAS^V12^ cells +/-Dox and treated or not with 50 U/ml IFN-β. Mean values are shown for three independent experiments. (**E**) Number of 53BP1 foci in BJ-RAS^V12^ cells (+/-Dox) treated or not with 50 or 100 U/ml IFN-β. Mean values are shown for three independent experiments.

Next, we monitored the impact of IFN-β on RS. Using DNA fiber spreading, we observed a sharp reduction in replication fork velocity after IFN-β treatment (**Fig. 6C**), which was even stronger than RAS^V12^ induction. We also observed a concomitant increase in the frequency of micronuclei, which was additive with RAS^V12^ induction (**Fig. 6D** and **EV Fig. 6C**). IFN-β also increased the frequency of 53BP1 foci (**Fig. 6E**) and further enhanced their formation upon RAS^V12^ induction (**Fig. 6E** and **EV Fig. 6E**). Along the same line, we observed an increase in both the frequency of micronuclei and the number of 53BP1 foci after overexpression of TREX1-D18N (**EV Fig. 6F** and **6G)**. This increase was comparable to that resulting from RAS^V12^ induction and both effects were additive (**EV Fig. 6D** and **6E)**. We also found that 30 to 40% of micronuclei colocalized with cGAS in untreated cells, which is consistent with earlier reports (Harding *et al*, 2017; Mohr *et al*, 2021; Mackenzie *et al*, 2017). Interestingly, the fraction of cGAS-positive micronuclei remained the same after IFN-β treatment (**EV Fig. 6D**) or overexpression of TREX1 D18N (**EV Fig. 6H**), regardless of the total number of micronuclei. However, it increased to ∼60% upon RAS^V12^ induction (**EV Fig. 6D**). Taken together, these results indicate that cells directly exposed to IFN-β or activating the cGAS-STING pathway due to their incapacity to degrade cytosolic DNA exhibit features of oncogene-induced RS, such as slower replication forks, accumulation of 53BP1 foci and micronuclei, independently of RAS^V12^ induction.

## Discussion

It is now well established that OIS can be triggered by two distinct events, namely replication stress and IFN signaling (Bartkova et al., 2006; di Micco et al., 2006; Dou et al., 2017; Glück et al., 2017; Yang et al., 2017), but whether these two pathways promote OIS independently or are functionally linked is currently unknown. The results presented in this study indicate that the nucleases MRE11 and TREX1 regulate both processes to promote senescence in BJ-RAS^V12^ cells, providing a mechanistic link between oncogene-induced RS and IFN signaling in OIS.

The MRE11 nuclease forms a heterotrimeric complex with RAD50 and NBS1 and plays pleiotropic roles in the maintenance of genome stability, including HR-mediated DSB repair, ATM activation and fork repair (Stracker & Petrini, 2011; Costanzo, 2011). As such, the MRN complex represents a barrier to oncogene-driven neoplasia (Gupta *et al*, 2013; Fagan-Solis *et al*, 2020). Importantly, MRE11 is also required to activate cGAS under RS conditions (Coquel *et al*, 2018; Pasero & Vindigni, 2017) and to induce SASP in senescent cells (Malaquin et al., 2020, Vizioli et al., 2020). Thus, MRE11 is strategically placed at the interface between oncogene-induced RS and IFN signaling. However, it is not known whether MRE11 links these two pathways in OIS. Here, we show that MRE11 is required for the slowing of replication forks, the formation of micronuclei, the induction of IFN, ISG and SASP genes, the growth inhibition and the induction of SA-β-gal activity in immortalized human fibroblasts overexpressing the H-RAS^V12^ oncogene. It plays therefore a central role in OIS by linking the RS and IFN responses to H-RAS^V12^ induction.

One of the earliest manifestations of RAS^V12^-induced RS is a dramatic reduction in fork velocity (Bianco *et al*, 2019; Aird *et al*, 2013; Di Micco *et al*, 2006). Our data show that this slow fork phenotype is suppressed by Mirin, suggesting that the tumor suppressive function of MRE11 comes at the cost of a slower replication. This fork slowing may represent a general fork protection mechanism, as MRE11 also reduces fork speed in cells expressing progerin (Kreienkamp *et al*, 2018), lacking PDS5 (Carvajal-Maldonado *et al*, 2019), or exposed to camptothecin (Rainey *et al*, 2020). The mechanism by which MRE11 activity regulates fork speed is currently unknown. It may be related to the formation of post-replicative ssDNA gaps (Hashimoto *et al*, 2010; Kolinjivadi *et al*, 2017), which activate ATR and may induce a checkpoint-dependent slowdown of DNA synthesis (Seiler *et al*, 2007; Mutreja *et al*, 2018; Frattini *et al*, 2021). Alternatively, MRE11 could modulate the formation or the stability of reversed forks (Lemacon *et al*, 2017; Pasero & Vindigni, 2017; Quinet *et al*, 2020), which could indirectly affect the speed of replication forks.

We recently reported that BJ-RAS^V12^ cells escape OIS by overexpressing components of the fork protection complex (FPC) such as Claspin and Timeless (Bianco et al., 2019). Cells overexpressing these factors also exhibit normal fork progression, suggesting that the FPC protects forks from RAS^V12^-induced RS. Here, we show that high levels of Claspin and Timeless also prevent the induction of IFN, ISG and SASP genes in BJ-RAS^V12^ cells. Taken together, our data support the view that oncogene-induced RS and IFN signaling are two sides of the same coin, potentially linked by MRE11. But how does MRE11 perform this function at the molecular level?

MRE11 exhibits both endonuclease and exonuclease activities, the latter being efficiently repressed by Mirin (Dupré *et al*, 2008). The onset of senescence hallmarks in BJ-RAS^V12^ cells was prevented by Mirin, suggesting that MRE11 acts through its exonuclease activity to promote OIS. However, we found that PFM01, a specific inhibitor of the endonuclease activity of MRE11, also restored the ability of BJ-RAS^V12^ fibroblasts to incorporate BrdU. It has been recently shown that PFM01 blocks fork breakage in absence of RAD51 (Mann *et al*, 2022). These data suggest that the cleavage of DNA strands could also contribute to OIS in BJ-RAS^V12^ cells, presumably through the release of cytosolic DNA. The presence of cytosolic ssDNA and dsDNA has been reported in senescent cells (de Cecco et al., 2019; Takahashi et al., 2018). Cytosolic DNA can be released by the processing of DNA ends and stalled replication forks by nucleases such as MRE11 and MUS81 (Lin and Pasero 2021; Coquel et al. Nature 2018; Erdal et al., 2017; Ho et al., 2016). Here, we were unable to detect increased levels of cytosolic DNA fragments upon RAS^V12^ induction. However, we observed higher micronuclei frequency in BJ-RASV12 cells another source of cytosolic DNA implicated in cGAS activation (Harding *et al*, 2017; Mackenzie *et al*, 2017; Mohr *et al*, 2021).

Micronuclei often form after mitosis around lagging chromosomes or chromosome fragments in cells exposed to chronic RS (Chan *et al*, 2009; Wilhelm *et al*, 2019). In BJ-RAS^V12^ cells, micronuclei formation was largely suppressed by Mirin, which is consistent with the view that micronuclei result from oncogene-induced RS. As reported by others (Harding *et al*, 2017; Mackenzie *et al*, 2017; Mohr *et al*, 2021; Glück *et al*, 2017; Yang *et al*, 2017), we observed an accumulation of cGAS in a subset of micronuclei in BJ-RAS^V12^ fibroblasts. We also confirmed that cGAS is involved in RAS^V12^-induced senescence (Dou et al., 2017; Glück et al., 2017; Yang et al., 2017). However, our data do not formally demonstrate that cGAS is activated inside micronuclei. Indeed, nucleosomes are known to sequester cGAS and prevent its activation (Pathare *et al*, 2020; Volkman *et al*, 2019; Kujirai *et al*, 2020). Determining how cGAS is activated upon RAS^V12^ induction and how this process is mediated by MRE11 will therefore require further work.

To investigate the contribution of cytosolic DNA and the cGAS-STING pathway in OIS, we overexpressed the major cytosolic nuclease TREX1 in BJ-RAS^V12^ fibroblasts. TREX1 counteracts cGAS signaling by degrading DNA in micronuclei (Mohr *et al*, 2021) and prevents the induction of SASP genes in senescent cells (Takahashi *et al*, 2018). Here, we found that TREX1 overexpression suppressed SA-β-gal activity in BJ-RAS^V12^ cells, supporting the view that cytosolic DNA activates cGAS in OIS.

More unexpectedly, we found that the overexpression of a dominant-negative form of TREX1 termed TREX1-D18N was sufficient to induce SA-β-gal activity in BJ cells to the same extent as RAS^V12^. Similarly, treatment of BJ cells with IFN-β inhibited BrdU incorporation in a dose-dependent manner. These results are intriguing because they suggest that the activation of the IFN pathway is sufficient to promote senescence independent of oncogene-induced RS. However, it has recently been reported that the ubiquitin-like protein ISG15, one of the major ISGs induced by type I IFNs, can also induce RS and DNA damage when overexpressed (Raso *et al*, 2020). These data suggest that the IFN response is not only induced by RS, but also acts as a positive feedback loop that further increases RS and OIS in response to RAS^V12^ induction. This view is supported by our data showing that long-term IFN-β treatment perturbs fork progression to the same extent as RAS^V12^ overexpression. In additions, both IFN-β and TREX1-D18N induced the formation of micronuclei and 53BP1 foci in BJ cells.

Altogether, these data argue for a model in which H-RAS^V12^ replication stress cross talks with the cytosolic DNA sensing pathway to promote senescence in a sophisticated manner. In concert with cytosolic nuclease TREX1, the nuclear nuclease MRE11 orchestrates RAS^V12^-induced senescence presumably through replication fork processing and cytosolic DNA accumulation. This work unveils the interplay of two distinct pathways of oncogene-induced senescence. Moreover, it brings new insights to the function of MRE11 and TREX1 in regulating cellular senescence.

Collectively, these data support a model in which the RS and cytosolic DNA sensing pathways do not act independently to induce OIS upon H-RAS^V12^ overexpression but are rather connected through MRE11 (**Fig. 7**). In this model, cytosolic DNA resulting from replication defects induces the formation of micronuclei and activates the cGAS-STING pathway. Importantly, the cGAS-dependent induction of IFNs and ISGs further increases RS, acting therefore like a positive feedback loop amplifying the RS response. This positive feedback loop explains why the addition of IFN-β is sufficient to induce RS, independently of H-RAS^V12^. This response to cytosolic DNA is placed under the control of the TREX1 nuclease, which prevents the unscheduled activation of OIS by degrading cytosolic DNA in normal cells.

**Figure 7.**
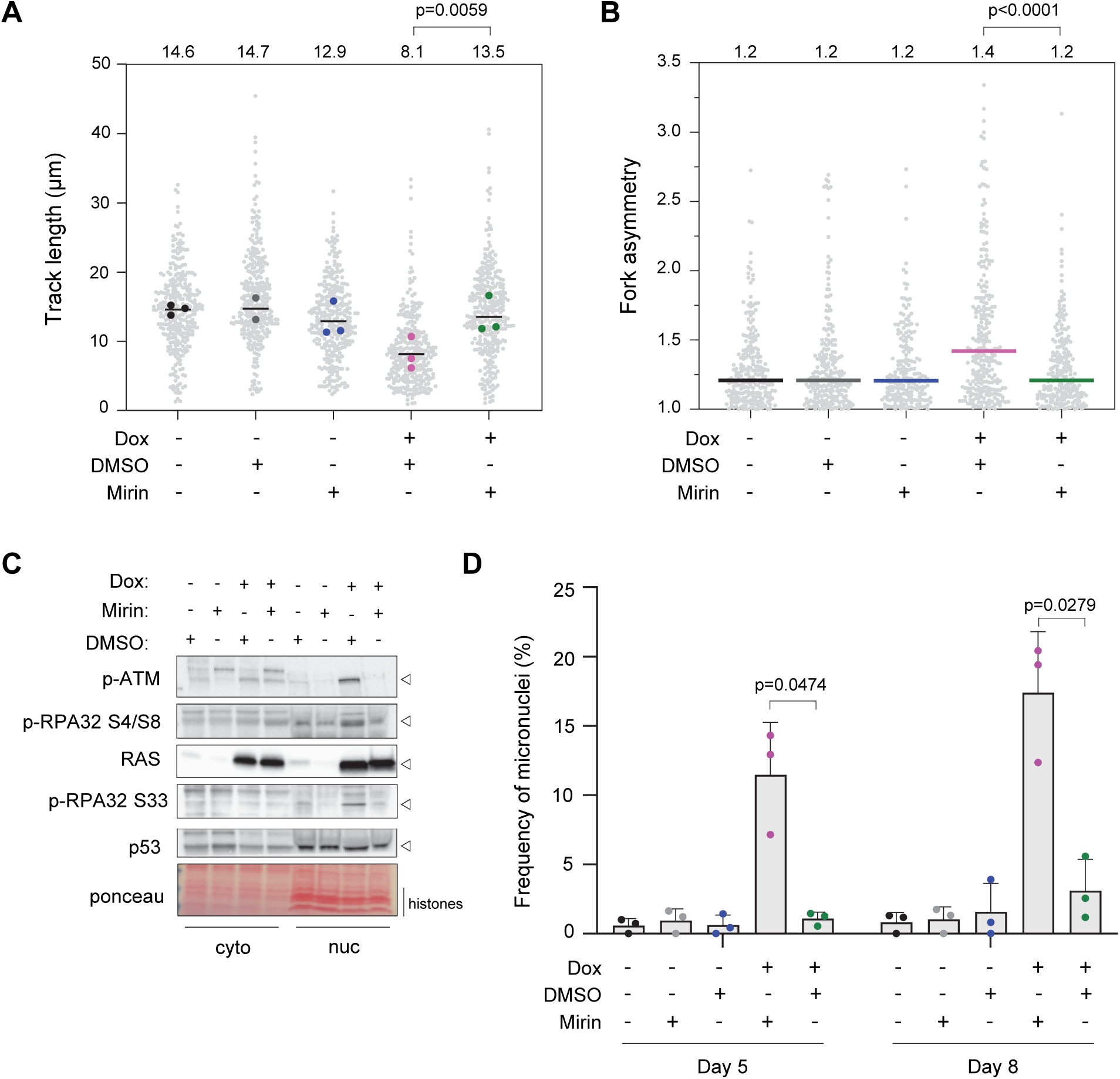
Model showing the connections between classical DDR and cytosolic DNA sensing during the establishment of senescence. (**A**) Classical model depicting two parallel pathways promoting senescence upon RAS^v12^ oncogenic stress. The DDR and cGAS-STING pathways both contribute to establish and maintain senescence. However, the links between these pathways and the exact contribution of cytosolic DNA to senescence were not known. (**B**) New model in which MRE11 links RS to IFN induction via the formation of micronuclei and cytosolic DNA in fibroblasts expressing the RAS^v12^ oncogene. This process is also controlled by the TREX1 exonuclease. Indeed, TREX1 overexpression prevents OIS and TREX1 inactivation in cells expressing the D18N dominant negative mutant promotes both genomic instability and senescence, even in absence of RAS^v12^. We propose that this positive feedback loop amplifies the signals that promote senescence by generating cytosolic DNA upon RS, activating IFN-signaling and further increasing RS and DDR activation.

## Materials and methods

### Cell culture

Normal human fibroblasts BJ-hTERT were a gift of Dr. D. Peeper (The Netherlands Cancer Institute, Amsterdam). H-RAS^V12^ from pBABE-Puro H-RAS^V12^ (n°12545 Addgene) was cloned in a home-made inducible vector derived from pLVX-Tight-Puro (Clontech) to make it SIN by a deletion in the 3’ LTR. Specifically, the XhoI-KpnI fragment of pLVX-Tight-Puro containing the inducible promoter was subcloned in pHR-BX (Lin *et al*, 2005) and the H-RAS^V12^ CDS was introduced into the MCS by appropriate restriction/ligation, as described previous (Bianco *et al*, 2019). BJ fibroblasts were grown in high glucose Dulbecco’s modified Eagle’s medium with ultraglutamine (DMEM), supplemented with 10% tetracycline-free and heat inactivated fetal bovine serum (FBS) and 100 U/mL penicillin/streptomycin (Lonza). Cell lines were grown at 37 °C in a humidified atmosphere of 5% CO2 and were tested for absence of mycoplasma contamination.

### Construction and production of TREX1 and TREX1(D18N) lentiviral vectors

Plasmids GFP-TREX1 and GFP-TREX1 (D18N), a gift from Judy Lieberman (Addgene plasmid 27219 and 27220, (Yan *et al*, 2010)), were digested with AgeI and XbaI and the fusion genes subcloned with the appropriate adaptators in pHRTK, a home-made, cPPT-containing SIN lentiviral vector derived from pHR-BX lentiviral vector (Lin *et al*, 2002) under the control of the constitutive CMV promotor. Virions were produced in HEK293T cells as previously described (Lin *et al*, 2004).

### Drugs and treatments

Doxycycline (Dox, purchased from SIGMA) was used at a final concentration 10 μg/ml (or otherwise stated in figure legends) to induce the RAS^V12^ overexpression in BJ-hTert fibroblasts. Mirin, PFM01 and PFM39 (all from SIGMA) were dissolved in DMSO and used at a final concentration of 10 μM. The cGAS (RU.521) and STING (H151) inhibitors were dissolved in DMSO and were purchased from Invivogen. According to drug treatments, the DMSO vehicle was used as a control. Recombinant human interferon-β was purchased from PeproTech (reference 300-02BC), dissolved in water at a stock concentration of 2 × 10^5^ Units per ml (U/ml) and stored in aliquots at -20°C. Recombinant IFN-β was used at final concentrations ranging from 50 to 300 U/ml as previously described (Glück et al, 2017).

### Western blot and antibodies

Cells were lysed in 2× Laemmli buffer at the concentration of 1 × 10^4^ cells/µl. Lysates were treated with 3 μl of Benzonase (25 U/µl, Sigma) for 30 min at 37 °C. Proteins were separated by sodium dodecyl sulfate polyacrylamide gel electrophoresis (SDS-PAGE), transferred to nitrocellulose membrane and analyzed by western immunoblotting with indicated antibodies: anti-RAS (1/500, BD, ref 610002), anti-MRE11 (1/500, Novus, NBS100-142), anti-cGAS (1/500, Cell Signaling, #15102), anti-p53 (1/500, Santa Cruz, clone DO7 sc-47698). Following antibodies were purchased from Abcam: anti-ATM (1:500, ab183324), anti-phospho S1981 ATM (1/500, ab81292), anti-IL1-a (1/500, ab9614), anti-ISG15 (1/1000, ab13346), anti-TREX1 (1:1000, ab185228). Rabbit anti-Phospho RPA32 (S33) and anti-Phospho RPA32 (S4/S8) were from Bethyl (used at 1/500, A300-246A and A300-245A, respectively). Blots were incubated with horseradish peroxidase-linked secondary antibody (GE Healthcare) and visualized using the ECL pico plus chemiluminescence method (Pierce) and a chemidoc camera (Biorad).

### SA-β-galactosidase staining

Cells were cultured in the presence of the indicated drugs and the indicated times. Then were processed with commercial SA-β-galactosidase staining kits following provider instructions (Kit #9860 from Cell signaling).

### Growth curve determination

At day 0, 3×10^5^ to 5×10^5^ of cells were seeded in 25cm^2^ flasks or in 6-well plates with the indicated treatments. At each passage (p), each 3 to 4 days, cells were counted with the automated cell counter - countess II (ThermoFisher) and all re-platted at the same density.

### DNA fiber spreading

DNA fibers were obtained and analyzed as previously described (Bianco *et al*, 2019; Coquel *et al*, 2018). Briefly, ongoing replication forks were sequentially pulse-labeled with thymidine analogues, 20μM IdU and then 100 μM CldU, for 20 minutes each. Cells were harvested by trypsinization and lysed in spreading buffer (200 mM Tris-HCl pH 7.5, 50 mM EDTA, 0.5% SDS) on microscope slides. After fiber fixation in methanol/acetic acid (3:1 ratio) the DNA fibers, and replication tracks were detected by immunostaining, using the following combination of antibodies. Primary antibody mix: Mouse anti-BrdU to detect IdU (1/100, BD 347580), Rat anti-BrdU to detect CldU (1/100, Eurobio clone BU1/75). Secondary antibody mix: Goat anti-rat Alexa 488 (1/100 PBS/T, Molecular Probes, A11006); Goat anti-mouse IgG1Alexa 546 (1/100 PBS/T, Molecular Probes, A21123). The ssDNA is detected with Anti-ssDNA (1/100, auto anti-ssDNA, DSHB) and then the goat anti-Mouse IgG2a Alexa 647 (1/50 PBS/T, Molecular Probes, A21241). Tracks were measured with FIJI. Statistical analysis of track lengths is performed with GraphPad Prism 9.0.

### DAPI staining to assess micronuclei frequency

To assess micronuclei formation, cells were grown on 12 mm diameter glass-coverslips (ref 0117520 Paul Marlenfeld GmbH & co), fixed 15-20 minutes in ice-cold 80% methanol, rinsed three times with PBS, and stained with a 1:5000 dilution of DAPI in PBS for 5-10 minutes at RT (DAPI was purchased from SIGMA). Alternatively, cells were fixed with 2% PFA 10 minutes at RT and then permeabilized with 0.1% Triton in PBS 10 minutes at RT. This last condition was used to estimate the frequency of cGAS positive micronuclei. Fixation was performed with 2% BSA in PBS 0.1% Triton for 45 minutes at RT. Immunostaining was performed overnight at 4°C with rabbit anti-cGAS D1D3G (1:400, Cell Signaling, #15102) and secondary goat anti-rabbit coupled to Alexa 546 (1:500, Molecular Probes, A11035). To estimate TREX1 localization to micronuclei we took advantage of the fluorescence signal produced by the expression of the tagged GFP-TREX1 proteins. After DAPI staining, coverslips were washed three times with PBS and mounted with ProLong Gold Antifade (P36930, ThermoFischer Scientific). Images were acquired with a Zeiss Axioimager Apotome microscope.

### BrdU staining

To assess the proliferation of BJ fibroblasts, cells were grown on glass coverslips (as described above) and then were pulse-labelled for 20 minutes with 30 μM of BrdU, then rinsed with PBS and fixed in ice-cold 80% methanol (as described above). Coverslips were then rinsed three times with PBS and stored at 4°C before immunostaining. DNA was denatured 45 minutes at RT in 2.5 M HCl, rinsed three times in PBS and then blocked 30-45 minutes in 2% BSA in PBS 0.1% Triton. For BrdU staining we used either a rat anti-BrdU (1:1000, Eurobio clone BU1/75) or mouse anti-BrdU (1:1000, BD 347580) antibodies 2 hours at RT (1:1000, Eurobio clone BU1/75). After 3 washes, fluorescently-conjugated secondary antibodies were used at 1:500 final dilution. Cells were DAPI stained, coverslips mounted in Prolong Gold Antifade (P36930, ThermoFischer Scientific) and images acquired with a Zeiss Axioimager Apotome microscope.

## 53BP1 foci staining

Cells were grown on glass coverslips as described above, then fixed with 2% PFA 10 minutes at RT and then permeabilized with 0.1% Triton in PBS 10 minutes at RT. Fixation was performed with 2% BSA in PBS 0.1% Triton for 45 minutes at RT. Immunostaining was performed overnight at 4°C with rabbit anti-53BP1 antibody (1:2000, Novus, NB100-304). After three washes, fluorescently conjugated antibodies were used to detect the signals (1:1000).

### RNA-seq

RNA sequencing (RNA-seq) were performed as previously described (Promonet *et al*, 2020; Bianco *et al*, 2019). Briefly, libraries were prepared using the Illumina TruSeq Stranded mRNA Library Prep Kit. Paired-end RNA-seq were performed with an Illumina NextSeq sequencing instrument (MGX-Biocampus, Montpellier, France). We focused our analysis on specific list of genes as follows. The hallmark of the Interferon-α response (IFN-α, from GSEA: M5911), Type I IFN-regulated genes with ISRE promoter elements (ISGs, from Reactome: R-HSA-9034125) and of SASP genes previously described (Ruiz-Vega *et al*, 2020).

### RT-qPCR

The expression of SASP markers IL6 and CXCL1 was quantified by qRT–PCR as described (Coquel *et al*, 2018) and normalized to *GAPDH*. Reverse transcription was performed by using the Superscript III First Synthesis System for RT–PCR (ref. 18080-051, Invitrogen). A LightCycler 480 SYBR Green I Master Mix (ref. 04887352001, Invitrogen) was used to perform quantitative PCR.

### Statistical analysis

All figures and statistical analysis were performed with GraphPad Prism 9.0. P-values of 0.05 or less were considered significant. The tests used are specified in the figure legends.

## Author contributions

H.T and YL.L, performed most of the experiments. H.T, wrote the original draft -edited and reviewed the manuscript and edited the figures. A.V, B.L and D.G provided technical assistance and contributed to image analysis. C.M constructed the TREX1, TREX1-D18N lentiviral vectors. D.G, J.H and J.M performed RNA-seq bioinformatics analysis. YL.L and P.P, conceived and supervised the study – edited and reviewed the manuscript and figures. All authors reviewed and accepted the manuscript before submission.

## Acknowledgements

H.T has been supported by AIRC under the iCARE-2 fellowship program. Our project has received funding from AIRC and from the European Union’s Horizon 2020 research and innovation program under the Marie Skłodowska-Curie grant agreement No 800924. Research carried by YL. L is supported by the Fondation ARC pour la recherche sur le cancer (N°PJA 20191209522). Work in the Pasero lab is supported by grants from the Agence Nationale pour la Recherche (ANR), Institut National du Cancer (INCa) and the Ligue Nationale Contre le Cancer (équipe labéllisée). We thank the MRI imaging facility (Biocampus) for assistance with image acquisition and analysis.

## Data Availability

The data sets generated and/or analyzed during the current study are available from the corresponding authors on reasonable request. The RNA-seq datasets showed in Figure 1 has been previously generated and reported (Bianco *et al*, 2019), and are available in the GEO repository, accession number: GSE123380.

## Figure legends

## Expanded View Figure legends

**EV Figure 1.**
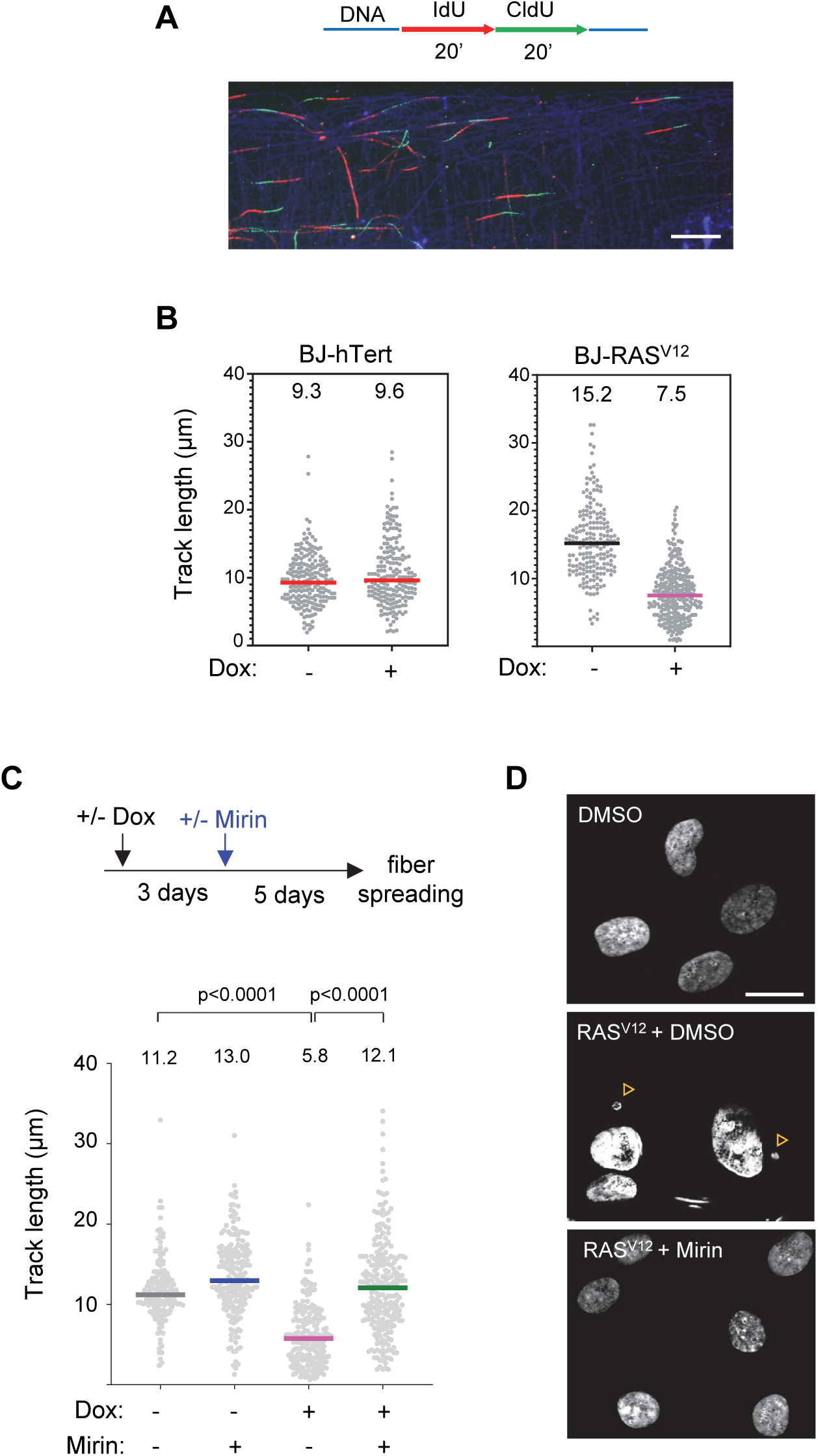
Volcano plots of differentially expressed IFN-α, ISG and SASP genes in senescent BJ-RAS^v12^ fibroblasts and in a BJ-RAS^v12^ clone escaping senescence independently of Claspin and Timeless overexpression (clone #8; Bianco *et al*, 2019). RNA-seq data (biological duplicates) were processed as indicated in Fig.1A.

**EV Figure 2.**
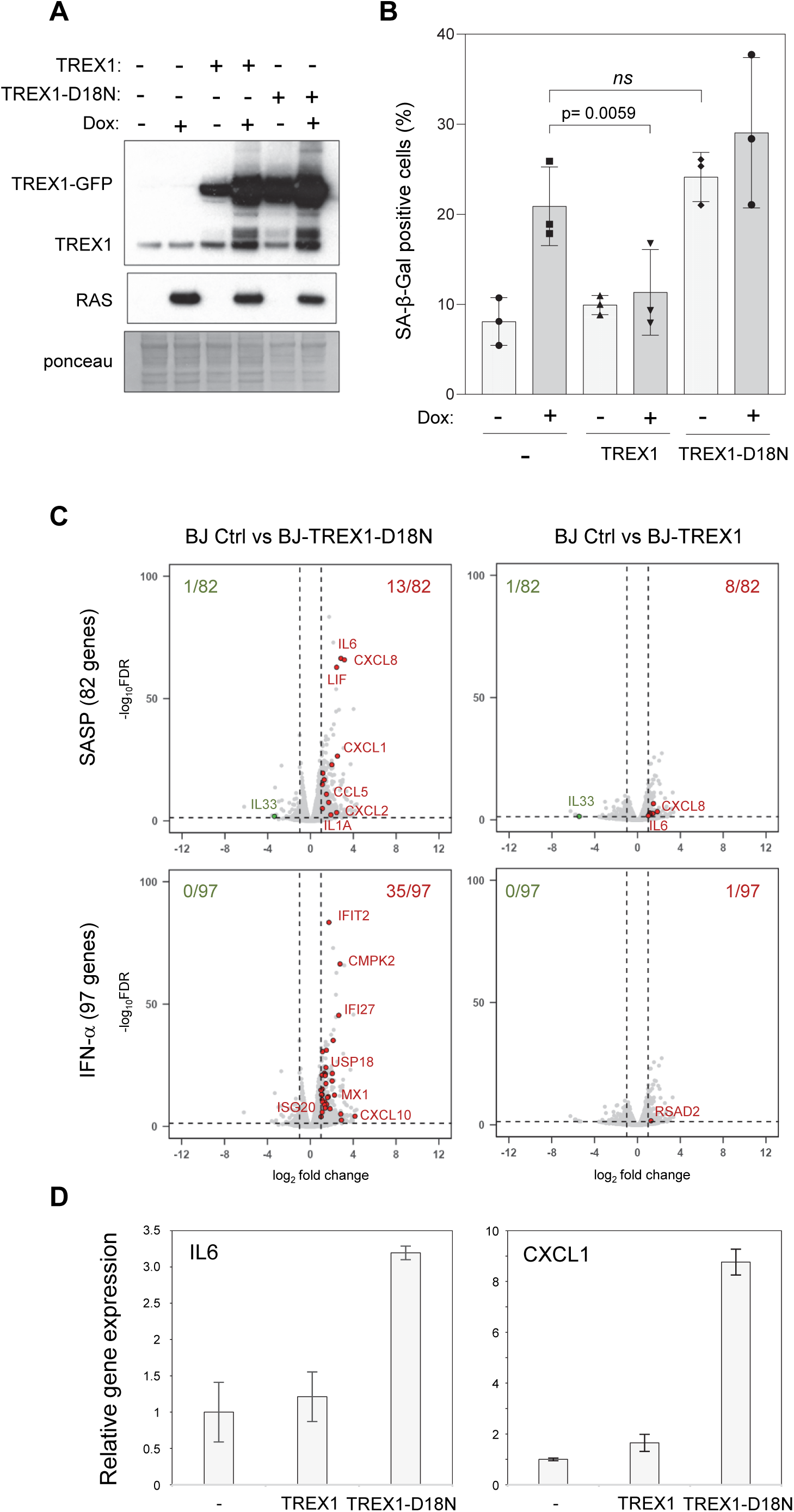
Western blot analysis of the expression of the SASP factor IL1-α in BJ cells overexpressing (+Dox) or not (-Dox) RAS^v12^ and treated or not with 10 µM Mirin. Cytosolic and nuclear fractions were obtained by lysing cells for 5 minutes on ice with a lysis buffer (10 mM Tris pH7.4, 10 mM NaCl, 10 mM MgCl_2_) containing 0.25% NP-40. Nuclear pellets were lysed in 2x Laemmli buffer and digested with benzonase. GAPDH and ponceau staining are shown as fractionation and loading controls.

**EV Figure 3.**
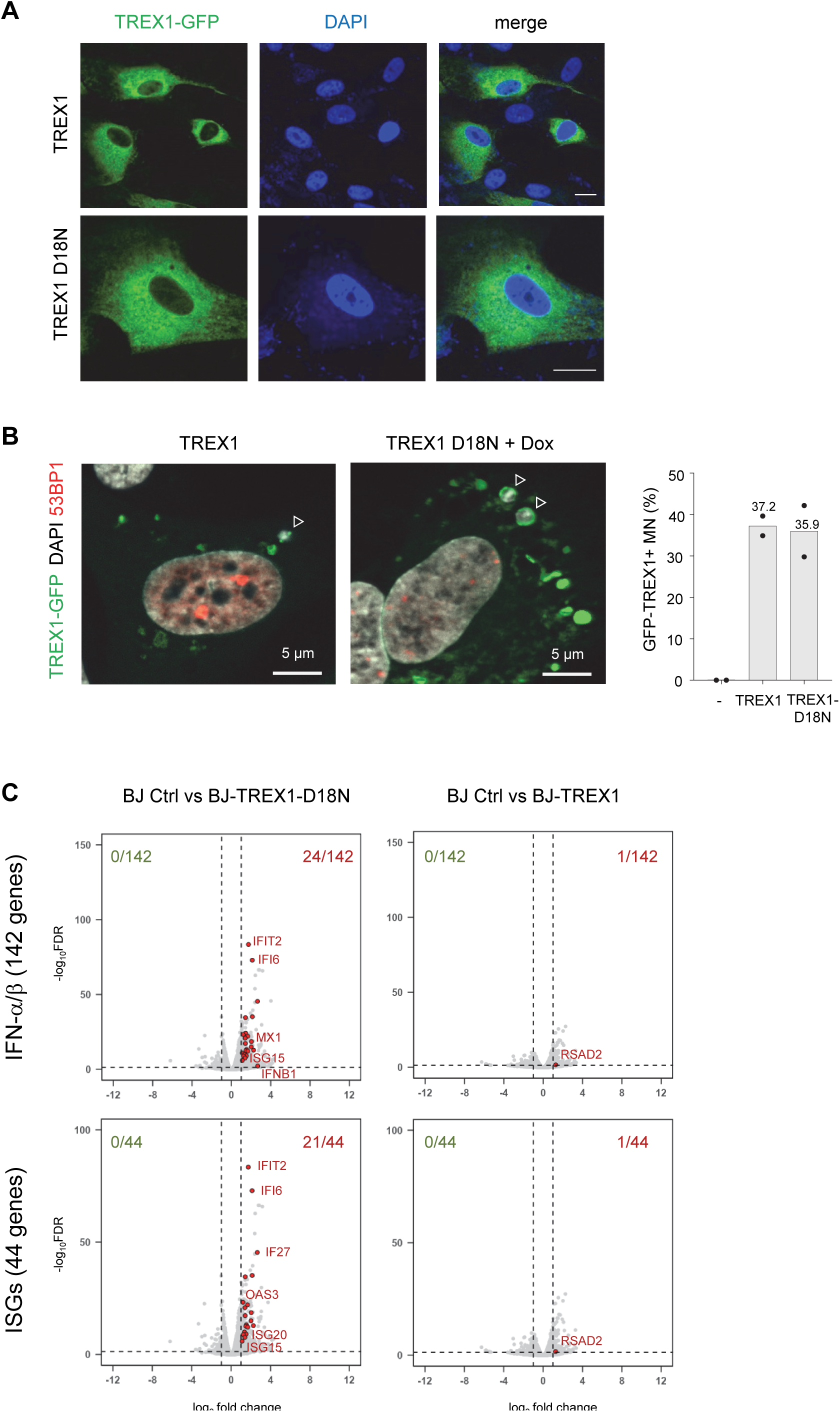
Mirin suppress OIS in BJ-RAS fibroblasts. (**A**) Representative pictures of SA-β-gal staining in BJ-RAS^V12^ fibroblasts overexpressing RAS^v12^ and treated or not with 10 µM Mirin or DMSO for 8 and 14 days. Scale bars are 100 μm. (**B**) Proliferation assay was performed by BrdU staining. Mirin treatment was performed five days post RAS^v12^ induction (+ Dox) for three days. DAPI is in blue, and BrdU is in green. Scale bars are 100 μm. The percentage of BrdU positive-cells is indicated for each condition (n=3).

**EV Figure 4.**
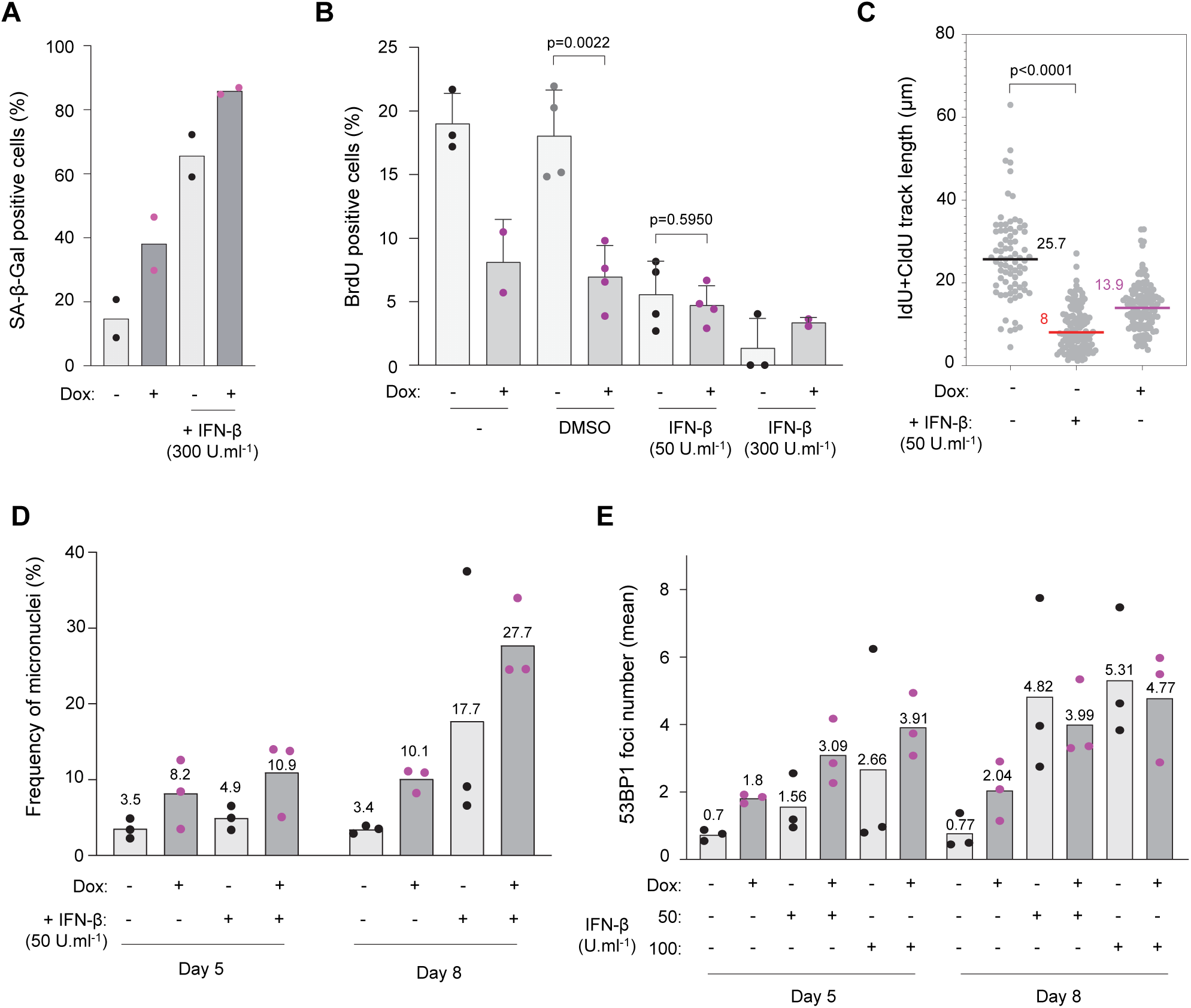
Mirin suppress replication stress in BJ-RAS^V12^ fibroblasts. (**A**) Experimental scheme. Cells were treated with Dox or Mirin for 8 consecutive days and fork progression was analyzed by DNA fiber spreading. Cells were pulse-labeled with two consecutive pulses of IdU and CldU for 20 minutes each. DNA was counterstained with an anti-ssDNA antibody (blue) and monoclonal antibodies against IdU and CldU (red and green, respectively, see Materials and Methods for details). Ongoing forks appear as adjacent red and green tracks. Total track length is measured as the sum of individual red and green tracks. (**B**) Fork progression in BJ-hTert (n=3) and BJ-RAS^v12^ cells (n=5) treated or not with Dox. BJ-hTert cells are used here as a control of the effect of Dox alone on fork speed. (**C**) DNA fiber analysis of fork progression in BJ-RAS^V12^ cells treated or not with Dox for 8 days and treated or not with 10 µM Mirin for the last 3 days of Dox induction. Results are representative of two independent experiments. (**D**) Representative images of DAPI staining in BJ-RAS after induction of RAS^v12^ for 5 days (Dox at 10 μg/ml) and treated with Mirin or DMSO. Arrow heads point to micronuclei. Scale bar is 20 μm.

**EV Figure 5.**
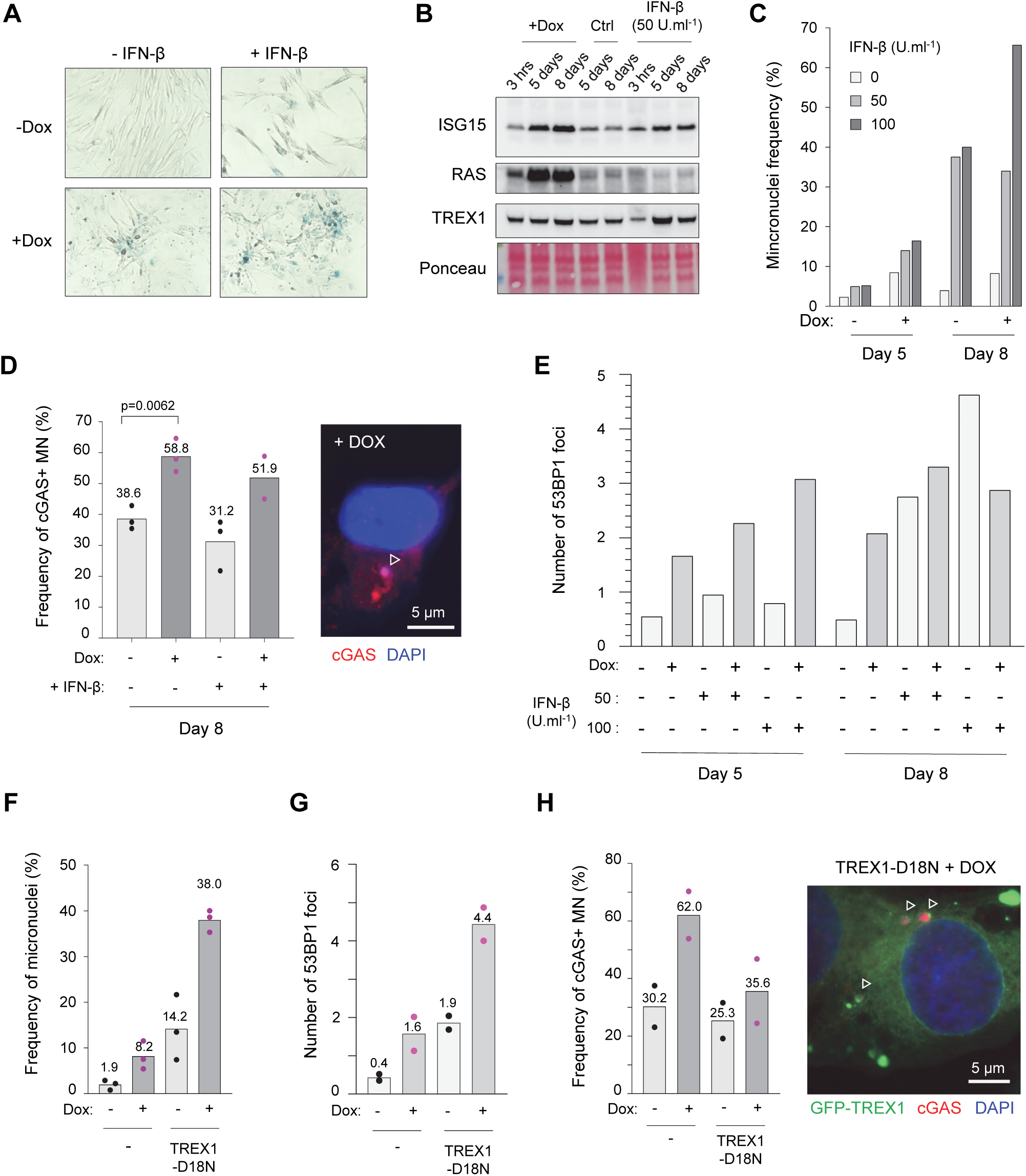
TREX1 regulates the IFN response in BJ-RAS fibroblasts. (**A**) Subcellular localization of TREX1-GFP and TREX1-D18N-GFP. Images are representative of at least three independent experiments. (**B**) TREX1-GFP colocalizes with micronuclei. The frequency of TREX1-GFP positive micronuclei (TREX1-GFP^+^ MN) is shown for two independent experiments. Representative images are shown. (**C**) Volcano plots of differentially expressed IFN-α/β (Reactome) and ISG genes in control BJ cells and in cells overexpressing either TREX1 or TREX1-D18N. RNA-seq samples were processed as indicated in Fig. 5C.

**EV Figure 6.**
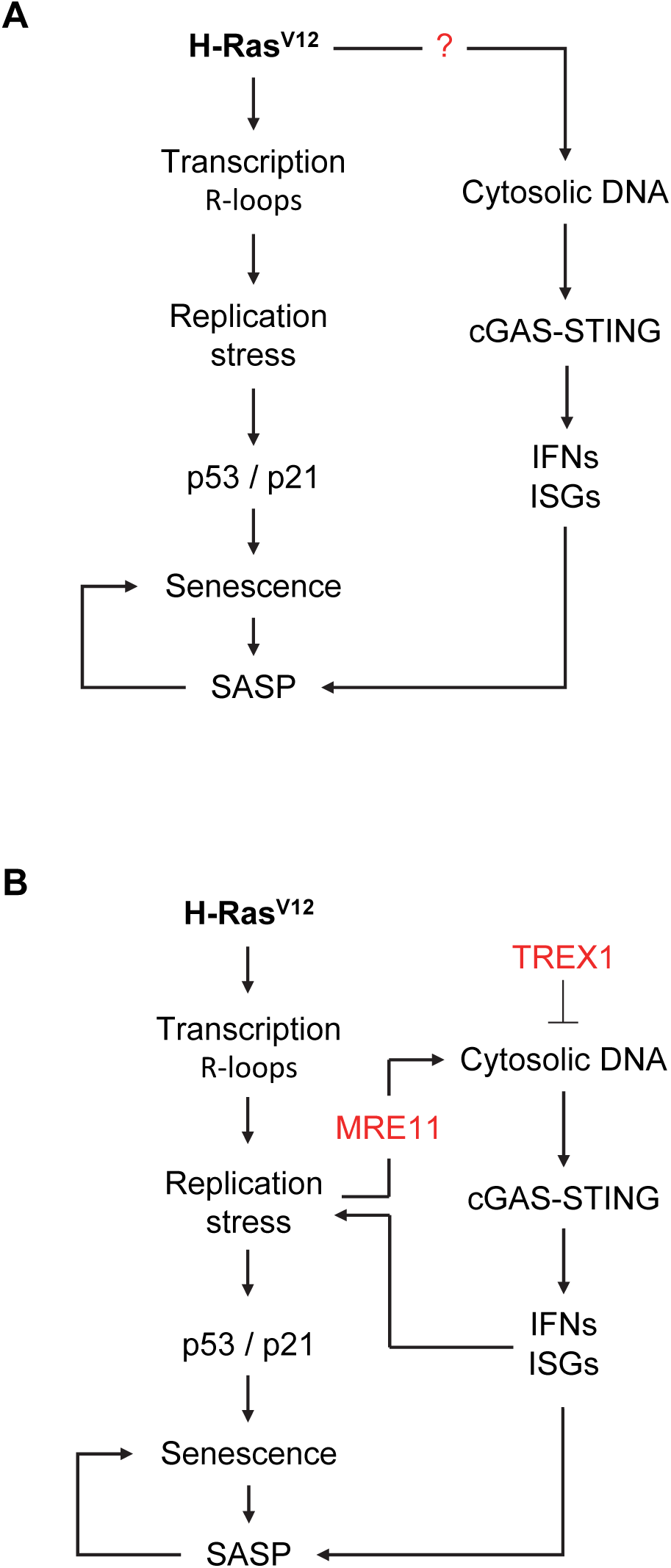
IFN-β induces DNA breaks and micronuclei in BJ cells. (**A**) Representative images of SA-β-gal staining in cells shown in Fig. 6A. (**B**) Western blot analysis of ISG15 induction by recombinant IFN-β. Cells were treated for 3 hours, 5 days or 8 days with Dox or INF-β as indicated (n=2). (**C**) Frequency of micronuclei in cells supplemented with either 50 or 100 U/ml IFN-β for 5 or 8 days. A representative experiment is shown. Biological triplicates treated with 50 U/ml of IFN-β are shown in Fig. 6D. (**D**) Immunofluorescence analysis of the localization of cGAS to micronuclei. Mean values are shown for at least two independent experiments. Representative image showing a cGAS-positive micronucleus (arrowhead). Scale bar is 5 μm. Right panel. (**E**) Number of 53BP1 foci in BJ-RAS cells treated with Dox and IFN-β (50 or 100 U/ml). Biological triplicates are shown in Fig. 6E. (**F**) Frequency of micronuclei in cells overexpressing RAS^v12^ and the dominant negative mutant of TREX1 (TREX1-D18N; n=3). (**G**) Number of 53BP1 foci from cells overexpressing RAS^v12^ and the dominant negative mutant of TREX1 (TREX1-D18N; n=2). (**H**) Frequency of cGAS-positive micronuclei in BJ cells overexpressing RAS^v12^ and the TREX1-D18N mutant (n=2). The localization of cGAS was assessed by immunofluorescence. A representative image showing cGAS-positive micronuclei (arrowheads) is shown. Scale bar is 5 μm.

